# The ubiquitous terpene geosmin is a warning chemical

**DOI:** 10.1101/2021.03.09.434661

**Authors:** Liana Zaroubi, Imge Ozugergin, Karina Mastronardi, Anic Imfeld, Chris Law, Yves Gélinas, Alisa Piekny, Brandon L. Findlay

**Affiliations:** Department of Chemistry and Biochemistry, Concordia University, Montreal, Québec, Canada; Department of Biology, Concordia University, Montreal, Québec, Canada

**Author notes:** Correspondence to: Brandon L. Findlay.

## Abstract

Known as the smell of earth after rain, geosmin is an odorous terpene detectable by humans at picomolar concentrations. Geosmin production is heavily conserved in actinobacteria, myxobacteria, cyanobacteria, and some fungi, but its biological activity is poorly understood. We theorized that geosmin was an aposematic signal used to indicate the unpalatability of toxin-producing microbes, discouraging predation by eukaryotes. Consistent with this hypothesis we found that geosmin and the related terpene 2-methylisoborneol reduced predation of *Streptomyces coelicolor* and *Myxococcus xanthus* by the bacteriophagous *Caenorhabditis elegans*. Predation was restored by the removal of both terpene biosynthetic pathways or deletion of the *C. elegans* ASE sensory neuron, and resulted in the death of the nematodes. Geosmin itself was non-toxic. This is the first warning chemical to be identified in bacteria or fungi, and suggests molecular signalling affects microbial predator-prey interactions in a manner similar to the well-studied visual markers of poisonous animal prey.

## Introduction

Few natural products are as widespread as geosmin, the smell of wet earth ^1^. Its chief producers in the soil, the saprophytic bacterial phylum *Actinobacteria* and predatory/saprophytic bacterial order *Myxococcales*, are found on every continent, including Antarctica ^2,3^. Geosmin is also a common contaminant in drinking water and farmed fish ^4,5^, produced by aquatic equivalents of these soil bacteria and by cyanobacteria ^4^. Though chiefly a prokaryotic metabolite, geosmin is also produced by a range of fungi ^6,7^, and is found in the peel of beets ^8^. Geosmin production is heavily conserved ^9-12^, and frequently accompanied by another odorous terpene, 2-methylisoborneol (Figure 1A) ^10,13^.

**Figure 1:**
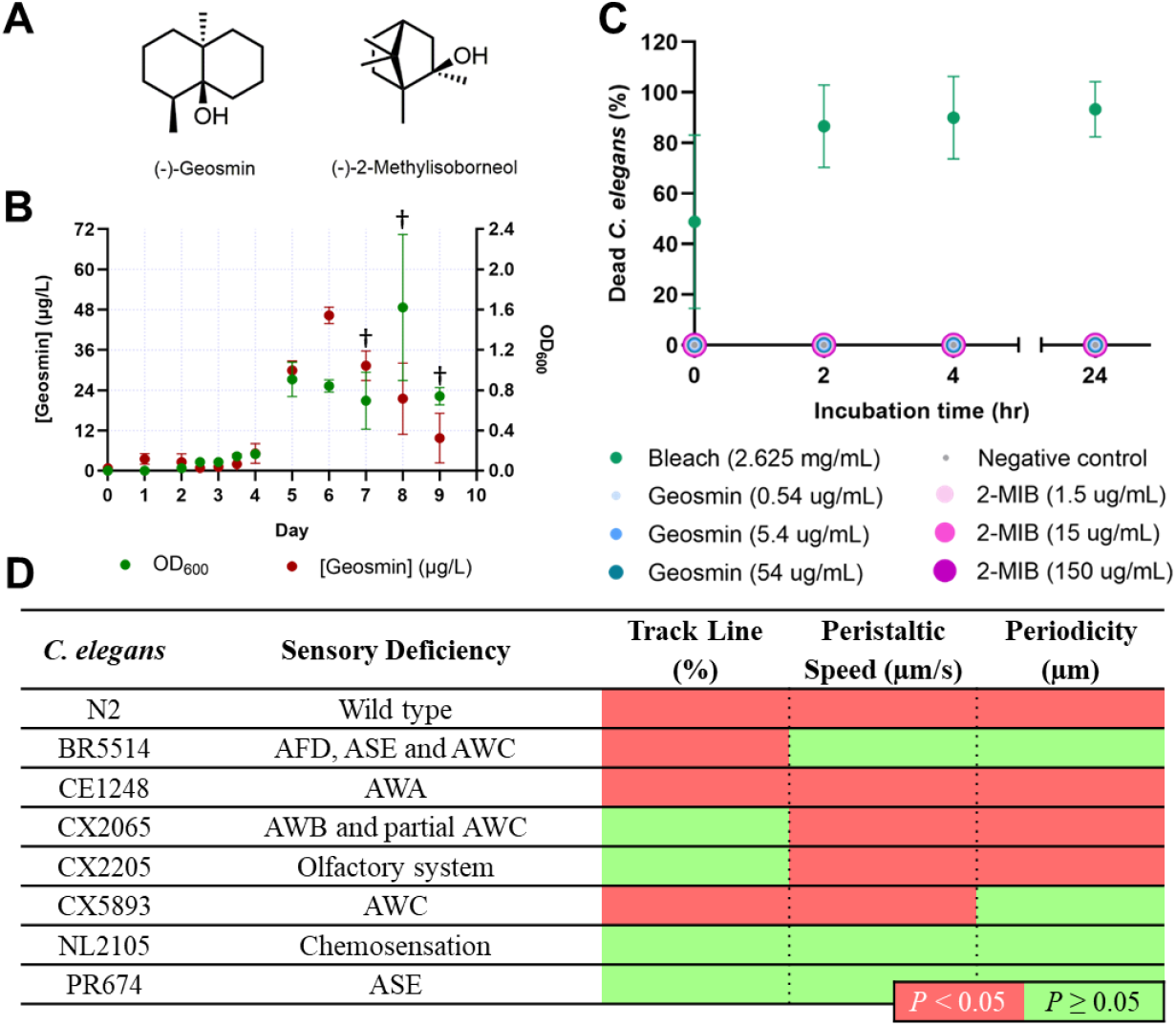
Geosmin production and toxicity. **A,** Chemical structures of geosmin and 2-methylisoborneol. **B,** Growth curve and geosmin production of *M. xanthus* DK1622 in 1% CTT. Geosmin concentrations were determined via GC-MS, with the aid of a calibration curve derived from solutions of authentic geosmin. † Cells began to clump at day 7, complicating OD_600_ measurements. **C,** Viability of adult *C. elegans* hermaphrodites in the presence of geosmin, 2-methylisoborneol and bleach. Worms non-responsive to touch stimuli were declared dead. Experiments were conducted in triplicate with five adult *C. elegans* per well (n=15). **D,** Comparison of the movement of adult *C. elegans* hermaphrodites on NGM and NGM-geosmin (0.54 μg/mL) plates, quantified with Imaris (track line) and WormLab (peristaltic speed, periodicity). Significant differences between NGM and NGM-geosmin plates are represented by a red box (*P* < 0.05) and non-significant differences by a green box (*P* ≥ 0.05). Experiments were conducted in triplicate with three worms per plate (n=9). Full details are available in External Data Table 4.

The breadth of geosmin producers and their varied ecological niches complicates assignment of the compound’s broader function. Geosmin repels egg-laying *Drosophila melanogaster*, which has a dedicated olfactory sensory neuron for geosmin detection ^14^, and attracts mosquitos and ants ^8,15^, Geosmin attracts springtails to sporulating *Streptomyces* colonies, which then eat the bacteria and help disperse bacterial spores ^16^, but this interaction would be of limited utility in aquatic environments ^12^. Due to differing primary nutrient sources, life cycles, cell wall structures, predators, and symbionts the principal geosmin producers have few features in common ^9,10,12^, but all produce a wealth of non-geosmin secondary metabolites ^17^. Many of these bioactive compounds inhibit the growth of prokaryotes or eukaryotes ^17,18^, and may be used in nature to deter competitors and predators ^19-22^.

In the animal kingdom, toxic prey advertise their unpalatability through the use of warning colours ^23^. These bright colours make the prey more conspicuous, but when combined with negative stimuli they deter predation through learned responses ^24,25^. To date no warning colours or other aposematic signals have been identified in prokaryotes, though olfactory signals may be used to reduce scavenging of nutrient-rich insects killed by entomopathogenic bacteria ^26^.

We propose that geosmin is a warning chemical, expressed by actinobacteria, myxobacteria, and cyanobacteria to advertise the production of toxic secondary metabolites. Commensurate with this hypothesis geosmin was non-toxic to the bacteriophagous nematode *Caenorhabditis elegans*, and did not prevent the worms from preying on non-toxinogenic *Escherichia coli* cells. Geosmin also did not act as a nematode repellent or attractant, but through interactions mediated by the ASE gustatory neuron triggered a distinct crawling behaviour with characteristics of both roaming and dwelling movement. Geosmin and 2-methylisoborneol significantly reduced nematode grazing on *Streptomyces coelicolor* colonies, though the terpenes were non-toxic to *C. elegans*. Grazing in the absence of the terpenes or by *C. elegans* mutants that lacked the ASE gustatory neuron triggered sporulation and the expression of a toxic secondary metabolite by *S. coelicolor*, while consumption of *S. coelicolor* or *M. xanthus* led to a *C. elegans* stress behaviour and the death of the worms. The use of geosmin as a warning chemical that deters predation by signalling the presence of toxic secondary metabolites is in line with previous reports of its attractant and repellant activities, and explains the high prevalence of geosmin biosynthetic genes in evolutionarily disparate microbes well known for their toxic natural products.

## Results

### Geosmin is produced during growth

To determine when geosmin provides the most benefit to its producers we characterized the production of geosmin as a function of growth in the predatory bacteria *M. xanthus* DK1622. In liquid media geosmin concentrations rose during the bacteria’s exponential phase and plateaued during early stationary phase, decreasing thereafter (Figure 1B). Fractionation of the cells and media and subsequent extraction with ethyl acetate confirmed that the majority of geosmin was excreted from the cell (Table S1). This production of geosmin during exponential phase suggests the compound is relevant when *M. xanthus* is hunting other bacteria.

### Geosmin does not aid protein degradation

Myxobacteria and actinobacteria obtain nutrients through the degradation of organic matter, either as saprophytes or predators ^2,27^. After confirming that geosmin did not inhibit the growth of Gram-negative or Gram-positive bacteria at physiologically relevant concentrations (Table S2), we examined the effect of geosmin on protease activity. Geosmin slightly reduced degradation of FTC-casein by the digestive enzymes excreted by *M. xanthus* DK1622 ^28^, and had no effect on degradation by a well-characterized human protease, trypsin (Figure S1).

As geosmin associates with membrane proteins in cyanobacteria ^4^, we postulated that it might stabilize hydrophobic digestive enzymes in *M. xanthus*. To evaluate this function we measured the effect of geosmin on heat-induced denaturation of the model hydrophobic protein bovine serum albumin (BSA). Geosmin had no discernable effect on BSA stability as measured by circular dichroism, either at elevated temperatures or on heating or cooling (Figure S2).

### Geosmin alters nematode behaviour

To determine if geosmin inhibited *M. xanthus* predators we tested its toxicity against the nematode *C. elegans*. In addition to their key role in understanding the development and function of the nervous system ^29^, nematodes are prevalent in soils across the globe and their grazing impacts microbial abundance and diversity ^30,31^. When mixed into agar geosmin had no effect on the viability of adult hermaphrodite nematodes over a 24-hour period (Figure 1C, Figure S3A), and did not appear to act as a chemoattractant or chemorepellent (Table S3). Grazing on the standard laboratory *C. elegans* prey, *E. coli* OP50, was similarly unaffected by geosmin, even at terpene concentrations well above that produced by *M. xanthus*, as evidenced by deep tracks in the *E. coli* lawn after 24 hr of incubation (Figure S3B-C).

However, geosmin strongly altered the worms’ movement. On NGM agar plates adult hermaphrodites exhibited clear roaming behaviour ^32^, with long, slightly arching pathways and few distinct turns or reversals. On NGM-geosmin plates the nematodes immediately began to move erratically, with many abrupt turns. Unlike traditional dwelling behaviour or responses to other bacterial metabolites ^32,33^, these worms moved faster than control worms and did not frequently reverse direction (Table S4, Files S1-S2). To quantify this change in behaviour we followed the movement of the worms in Imaris and WormLab, measuring significant changes in track linearity, peristaltic speed, and head movement periodicity (Figure 1D, Table S4).

To determine how geosmin was causing these changes we compared the movement of a range of *C. elegans* mutants in the presence or absence of geosmin. A mutant deficient in chemosensation, *C. elegans* NL2105, exhibit similar behaviour in the presence and absence of geosmin, indicating that the terpene’s effect was mediated by the host chemosensation machinery (Figure 1D, Files S3-S4) ^34^. The modified dwelling behaviour was retained in worms deficient in the sensing of volatile attractants and repellents (mutants lacking AWA and AWB neurons, Figure 1D), suggesting that geosmin is not detected by the nematode olfactory system, but lost in worms lacking the gustatory neuron ASE. ASE has been previously linked to the sensing of water-soluble attractants and movement away from food as prey populations decline ^35,36^.

### Geosmin deters feeding on its producers

As geosmin did not limit predation of *E. coli* by *C. elegans*, to test its effect *in situ* we added *C. elegans* to plates containing colonies of *S. coelicolor*, using both the wildtype *S. coelicolor* M145 and mutant strains lacking production of geosmin (J3003) and both geosmin and 2-methylisoborneol (J2192)^16,37^. When *C. elegans* N2 was added to *S. coelicolor* M145 or J3003 the majority of worms localized outside of the bacterial colonies after 4 hr (Figure 2A). When added to *S. coelicolor* J2192 worms were predominantly found within the bacterial colony at the 2 hr and 4 hr mark. Worms lacking the ASE neuron, *C. elegans* PR674, localized within all three bacterial strains at all time points (Figure 2B). In all experiments nematodes consumed the bacteria, as noted by the presence of red bacteria within the nematode pharynx (Figure 2E). The addition of worms lead to rapid sporulation of *S. coelicolor* and the production of the toxic bacterial metabolite actinorhodin (Figure 2G) ^38^. The majority of worms within bacterial colonies at the 24 hr mark were coated in white bacterial spores, and either exhibited distress behaviour (File S5), or appeared dead. The movement of worms into and out of bacterial colonies did disperse some bacterial spores (Figure 2G), but given the high toll on both bacterial growth and worm viability the overall effect of nematode predation was detrimental to both nematodes and bacteria.

**Figure 2:**
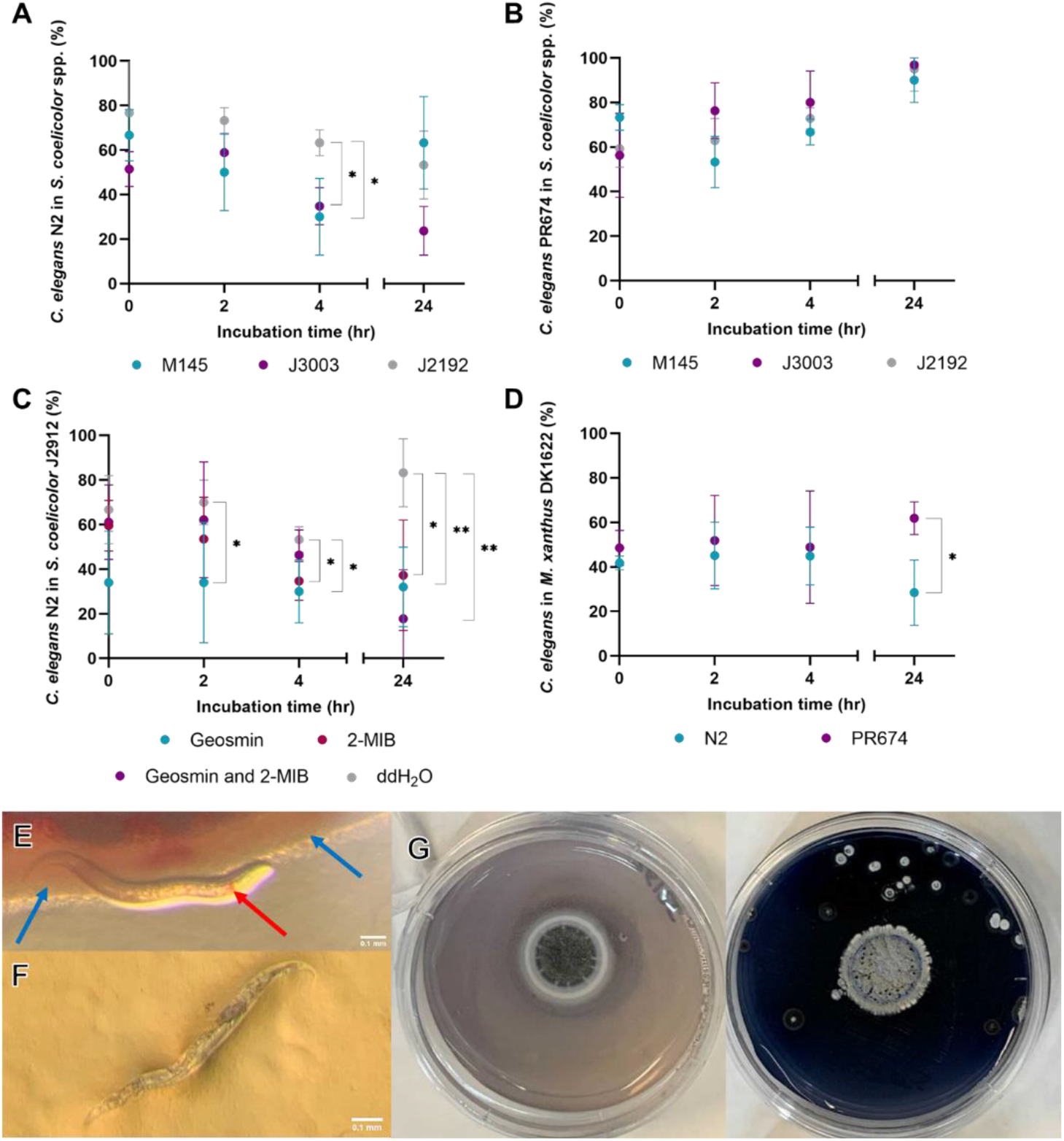
Predation of *S. coelicolor* and *M. xanthus* by *C. elegans*. **A,** Proportion of adult *C. elegans* N2 worms in colonies of *S. coelicolor* M145 (WT), J3003 (*ΔgeoA*) and J2192 (*ΔgeoA ΔmibAB*) as a function of time. **B,** Proportion of *C. elegans* PR674 (*che-1(p674)*, ASE deficient) worms in colonies of *S. coelicolor* M145 (WT), J3003 (*ΔgeoA*) and J2192 (*ΔgeoA ΔmibAB*) as a function of time. **C,** Proportion of *C. elegans* N2 worms in colonies of *S. coelicolor* J2912 that were pre-treated with geosmin, 2-methylisoborneol, or distilled, deionized water as a function of time. **D,** Proportion of *C. elegans* N2 and PR674 worms in colonies of *M. xanthus* DK1622 as a function of time. **E,** Consumption of *S. coelicolor* by *C. elegans*. Blue arrows indicate the bacterial colony, the red arrow indicates the presence of bacteria in the *C. elegans* pharynx. **F,** Production of spores and actinorhodin by *S. coelicolor* in the absence (left) or presence (right) of *C. elegans*. 10 day cultures, room temperature. **G,** *C. elegans* on a *M. xanthus* lawn. Worms became translucent prior to death and were ultimately digested by the bacteria. All nematode experiments were run in triplicate, with ten adult hermaphrodites per study (n=30). Worms were grown on *E. coli* OP50 and starved for 20 minutes prior to addition to *S. coelicolor* or *M. xanthus*. Statistically significant deviations from J2192 (A, B), H_2_O (C), or wildtype (D) are indicated by * (P < 0.05) or ** (P<0.01). Representative images are shown.

To ensure the non-terpene metabolite profile of *S. coelicolor* mutants J3003 and J2192 did not meaningfully differ from that of the wildtype we conducted a series of add-in experiments. Geosmin and 2-methylisoborneol were added alone or in combination to colonies of *S. coelicolor* J2192, at concentrations approximating their physiological values ^4,16^. Geosmin or 2-methylisoborneol was sufficient to significantly reduce the number of worms in bacterial colonies at the 2 hr, 4 hr, and 24 hr mark (Figure 2C). At 24 hrs few worms were found in bacterial colonies treated with geosmin and 2-methylisoborneol, in line with the co-occurrence of geosmin and 2-methylisoborneol biosynthetic genes in actinobacteria and cyanobacteria, but at the concentration of terpenes in our study the effect was not statistically distinct from 2-methylisoborneol or geosmin alone ^10,12^.

To determine if the reduction in predation that we observed was limited to *S. coelicolor* we repeated our assay with *M. xanthus* DK1622. As with the *Streptomyces, C. elegans* N2 localized outside the *M. xanthus* colony at the 24 hr mark, despite the latter’s swarming movement over the agar plate (Figure 2D). *C. elegans* PR674 showed no such aversion, resulting again in the death of both predator and bacterial prey (Figure 2F).

## Discussion

The high prevalence of geosmin synthase genes across unrelated bacteria and fungi suggests that geosmin is key to the fitness of a broad range of microorganisms. Here we propose that geosmin is a widespread warning chemical, used to advertise the toxicity of its producers and deter predation. Consistent with this role geosmin is produced during the mobile hunting phase of *M. xanthus* growth, and is actively excreted from the cell (Figure 1B, Table S1). Production in *Streptomyces* spp. is more complex, as the geosmin precursor isopentenyl pyrophosphate is produced by both the methylerythritol phosphate (MEP) and mevalonate pathways in this genus ^4^, and the two pathways are differentially regulated between strains and growth phases ^39,40^. Cyanobacteria stockpile geosmin and 2-methylisoborneol during exponential growth and stationary phase, and release the terpenes as they die ^41^.

To act as an aposematic signal geosmin must be detected by potential microbial predators. The bacteriophagous *C. elegans* detected geosmin through the gustatory neuron, ASE, switching to a behaviour characterized by frequent changes in direction and high movement speeds (Figure 1D, Table S4, Files S1-S4). In the presence of *S. coelicolor*, when worms were able to sense geosmin and either geosmin or 2-methylisoborneol was present this change in behaviour resulted in significantly fewer worms in the bacterial colonies (Figure 2A-B). Similar results were obtained when geosmin or 2-methylisoborneol was added to a *S. coelicolor* mutant deficient in the production of these terpenes, and when worms were added to geosmin-producing *M. xanthus* DK1622 (Figure 2C-D). Geosmin was not only non-toxic to these worms (Figure 1C), but by altering *C. elegans’s* feeding behaviour geosmin prevented the worms from coating their bodies in bacterial spores or ingesting toxic bacterial metabolites. As with Drosophila and *A. aegypti^8,14^*, this effect was mediated by the predator’s own chemosensory system.

The use of geosmin as an aposematic signal may explain its reported effects on other eukaryotes and its prevalence across a range of unrelated microbes. Geosmin attracts *Solenopsis invicta* because the terpene reliably indicates the presence of *Streptomyces* spp., and the toxic metabolites produced by these bacteria protect ant colonies from fungal infections ^15^. Similarly, geosmin discourages egg laying by Drosophila, whose young are susceptible to bacterial toxins ^14^, while also signalling the presence of edible cyanobacteria to the more toxin-resistant *A. aegypti* ^8,42^. In principle, acquisition of geosmin synthase by any toxin-producing microbe could recapitulate the aposematic phenotype, favouring lateral gene transfer between evolutionarily unrelated species and the evolution of Müllerian mimics ^43,44^. Other aposematic signals exhibit positive frequency-dependent selection ^45^, and the evolution of Müllerian mimics in bacteria likely favours the further lateral gene transfer of geosmin synthase. The ubiquity of geosmin in natural environments ensures few predators are naïve to the signal, while those that ignore it likely experience consistent fitness penalties from preying on toxic geosmin producers.

Geosmin did not prevent *C. elegans* from feeding on *E. coli* but heavily reduced grazing on *S. coelicolor* or *M. xanthus* (Figure 2), suggesting that aversion requires both the terpene and a negative stimuli ^33^. Microbial predators can discern between adjacent toxin-producing and non-toxic bacteria ^46^, and the ability of geosmin to deter but not protect against predation may prevent the emergence of Batesian mimics that express only the terpene. While *C. elegans* reacted to geosmin in the absence of *S. coelicolor* or *M. xanthus* (Figure 1D), it is unclear if the avoidance of toxin-producing bacteria was learned or innate. Higher eukaryotes can learn to associate aposematic signals with unpalatability ^24,47^, and while bacterial predators are significantly less complex, both *C. elegans* and amoeba can link sensory cues to past events ^48,49^.

In conclusion, geosmin is detected by the predatory nematode *C. elegans* through the gustatory neuron ASE. Geosmin is non-toxic to this species, but nematodes strongly avoid geosmin-producing bacteria. When geosmin production was eliminated or the ASE neuron was disabled *C. elegans* became coated in bacterial spores and ingested toxic secondary metabolites. Concurrently, bacterial fitness declined through nematode feeding and the conversion of vegetative cells to senescent spores. Geosmin thus acts as an aposematic signal, honestly and reliably advertising the unpalatability of its producers and providing a mutual benefit to predator and prey. Geosmin is the first warning chemical to be identified in bacteria, and it not only shapes bacterial predator-prey interactions but also appears to mediate interactions between eukaryotes and bacteria across the globe.

## Materials and Methods

### Strains and cultivation

#### Bacteria

*Klebsiella aerogenes* ATCC 13048 was acquired from the American Type Culture Collection (ATCC). *Escherichia coli* MG1655 and *Burkholderia thailandensis* E 264 were a gift from Eric Déziel, INRS-IAF. *Myxococcus xanthus* DK1622, *Micrococcus luteus* DSM 20030 and *Bacillus subtilis* DSM 10 were obtained from the Leibniz Institute DSMZ-German Collection of Microorganisms and Cell Cultures. *Streptomyces coelicolor* M145, J3003 and J2192 originated from the John Innes Centre, Norwich, UK, and were gifts from Dr. Klas Flärdh. *Escherichia coli* OP50 was obtained from the Caenorhabditis Genetics Center (CGC) which is funded by NIH Office of Research Infrastructure Programs (P40 OD010440).

*M. luteus, B. subtilis, B. thailandensis, K. aerogenes* and *E. coli* were grown in Luria-Bertani (LB) media, at 30 °C (37°C for *E. coli*) rotating at 225 rpm for liquid cultures. Bacterial isolates were streaked on 1.5% agar LB plates and placed at 30 °C for 24 hrs (37°C for 16 hrs for *E. coli*) prior to experiments. *M. xanthus* DK1622 was grown in 1% CTT at 30 °C and 225 rpm. Bacterial isolates were streaked on 1.5% agar CTT plates and placed at 30 °C for 3 days prior to experiments. *Streptomyces coelicolor* M145, J3003 and J2192 were grown at 30 °C in Tryptic Soy Broth (TSB) rotating at 225 rpm and isolates were streaked on 1.5% agar Tryptic Soy Agar (TSA). Growth curves for *M. xanthus* DK1622 were generated by performing daily OD_600_ measurements using a Varian Cary 100 Bio UV-vis spectrophotometer.

#### Caenorhabditis elegans

The *C. elegans* lineages were maintained on nematode growth medium (NGM) plates with *E. coli* OP50 at 20°C as per standard protocol ^50^. The wild type N2 and mutant strains BR5514 (*tax-2(p671);tax-4(p678)*), CE1258 (*eat-16(ep273)*), CX2065 (*odr-1(n1936)*), CX2205 (*odr-3(n2150)*), CX5893 (*kyIs140 I; ceh-36(ky646)*), NL2105 (*gpa-3(pk35) odr-3(n1605)*) and PR674 (*che-1(p674)*) were obtained from the Caenorhabditis Genetics Center (CGC) which is funded by NIH Office of Research Infrastructure Programs (P40 OD010440). Gravid nematodes were age synchronized and cleaned from bacterial and fungal contaminants using a bleaching mixture (2.5% NaClO, 0.5 M NaOH) before each experiment, as previously described ^50^. All *C. elegans* experiments were conducted using a standard stereomicroscope.

#### Chemicals

(±)-Geosmin standard was obtained from Sigma-Aldrich, (-)-geosmin was purchased from FUJIFILM WAKO chemicals. Bovine serum albumin, levamisole, (1R)-(+)-camphor and the 3.0 M methylmagnesium bromide solution in diethyl ether (189898) were obtained from Sigma-Aldrich. FTC-casein and trypsin proteins were both obtained from ThermoFisher Scientific. Methanol, ethanol, ethyl acetate, tetrahydrofuran (THF), hexane and 2-butanone were acquired from ACS Chemicals and Fisher Scientific.

#### Media and Buffers

1% CTT (1% casitone, 10 mM Tris-HCl [pH 7.6], 8 mM MgSO_4_, 1 mM KH_2_PO_4_) was prepared from scratch. LB (10 g/L tryptone, 10 g/L NaCl, 5 g/L yeast extract) was prepared from a premix, which was purchased from Bio Basic. TSB (17 g/L casein peptone, 3 g/L soya peptone, 5g/L NaCl, 2.5 g/L K_2_HPO_4_, glucose 2.5 g/L [pH 7.3]) premix was purchased from Sigma-Aldrich. NGM (3 g NaCl, 2.5 g peptone 20 g agar, 1 mL of 5 mg/mL cholesterol in ethanol, 1 mL of 1M MgSO_4_, 25 mL of 1M [pH 6.0] KPO_4_ in 1L H_2_O) was made from scratch. Worm M9 buffer (3 g/L KH_2_PO_4_, 6 g/L Na_2_HPO_4_, 5 g/L NaCl) was made from scratch. The TBS buffer used in the protease assays was made from scratch (25 mM tris, 0.15 M NaCl [pH 7.2]). The sodium phosphate buffer used in the CD assays as also prepared from scratch (pH 7.0, ionic strength 0.014 M).

#### Geosmin quantitation

A culture of *M. xanthus* DK1622 in stationary phase was diluted to an OD_600_ of 0.125, then diluted 1:100 in fresh 1% CTT. Samples were left shaking at 30 °C and 225 rpm. Aliquots were drawn every 12-24 hrs until day 9. OD_600_ measurements were made on a Varian Cary 100 Bio UV-vis spectrophotometer. Samples over an OD_600_ of 1.0 were diluted in fresh 1% CTT media to the 0.010-0.99 range. After 7 days incubation clumping was observed and prior to measurements, cells were dispersed via passage through a serological pipette. Geosmin extractions were made using a 1:1 ethyl acetate (EtOAc) extraction with *M. xanthus* DK1622 bacterial culture. Ethyl acetate samples were sonicated, then centrifuged for 1 min at 3000 g to collect clean supernatant before injection for quanitification by gas chromatography-mass spectrometry (GC-MS). To quantify media versus cytoplasmic geosmin the bacterial culture was centrifuged at 3000 g for 2 min. The supernatant was used to measure extracellular geosmin as detailed above, while the intracellular/cytoplasmic geosmin concentration was measured by exposing the pellet of *M. xanthus* cells to ethyl acetate, vortexing for 1 min, then sonicating and centrifuging down for 1 min at 3000 g. The GC-MS system (7890B GC coupled to a 5977B MS, Agilent Technologies) was equipped with an autosampler and a split/splitless inlet kept at 300°C. Splitless injections (1.0-3.0 μL) were made on a 60-m DB-EUPAH (0.25 mm ID x 25 μm film thickness; Agilent Technologies) column with the oven kept isothermal (80 °C) for 8 min, ramped to 300 °C at 15 °C /min, and then held at that temperature for 5 min. The inlet was kept at 300 °C throughout. Helium flow rate was 1.2 mL/min, with a He septum purge flow of 5 mL/min. Seven-level external calibration curves between 0.01 and 1.00 mg/L were used for quantitation.

#### Geosmin MIC

Following CLSI guidelines for direct colony suspension testing ^51^, bacterial cultures were transferred to LB broth and adjusted to a final turbidity equivalent to a 0.5 McFarland standard (1.5 × 10^8^ CFU/mL). Bacteria were then mixed 1:1 with (±)-geosmin in 96-well plates, then incubated at 30 °C for 20-24 hr. Methanol was used as a control. The MIC was defined as the concentration sufficient to inhibit bacterial growth as evaluated by the naked eye. After initial tests at concentrations similar to the level produced by *M. xanthus* DK1622 failed to inhibit growth the quantity was increased, until the methanol used to solubilize the high geosmin concentrations began to impede growth.

#### Effect of geosmin on BSA thermal denaturation

CD analyses were made using a Jasco J-715 Spectropolarimeter. The cuvette width was 0.2 cm. The BSA concentration was 1.25 μM for all experimental and control assays. Fresh protein samples were prepared before each experiment and kept on ice. The (-)-geosmin concentration was 0.18 mM for the experimental assay and the SDS concentration was 0.75 mM. All samples were diluted in PBS ^52,53^. Subtractions and smoothing were made using Jasco J-715 software using controls of each chemical in PBS exclusively. Elevated temperature analysis was made at 80°C. The wavelength window measured was from 200-240 nm. Five scans were made for each measurement. Experiments were run in triplicate before data analysis using the Jasco J-715 instrument. Varying temperature analysis was made at a wavelength of 208.6 nm and a heating/cooling rate of 1°C/min.

#### Protease activity assay

A Pierce™ Fluorescent Protease Assay Kit was purchased from Thermo Scientific™. The kit contained FTC-casein, n-tosyl-L-phenylalanine chloromethyl ketone (TPCK) treated trypsin standard and a tris-buffered saline (TBS) buffer pack ^54^.

As per the product sheets instructions ^54^, FTC-casein was diluted a 5 mg/mL stock solution with ultrapure water. One 20 μL aliquot of the stock solution was diluted 1:500 in TBS to a final volume of 10 mL and concentration of 0.01 mg/mL. A 20 μL aliquot of Trypsin stock solution (TSS) of 50 mg/mL was diluted to 1 mg/mL in TBS. To obtain the extracellular proteases of *M. xanthus* DK1622 absent geosmin the supernatant of a 10-day culture was extracted with EtOAc and the organic layer discarded. A 50% protease solution in TBS buffer was used and was kept in ice with FTC-casein and trypsin. Each fluorometric assay was measured using a Varian Cary Eclipse Fluorescence Spectrophotometer at λ_excitation_= 494 nm, and λ_emission_= 521 nm. The experiments were made with and without the presence of (±)-geosmin (1 mg/L) to evaluate the effect of (±)-geosmin on the activity of the proteases. Methanol was used as a negative control.

#### Chemotaxis assay

Chemotaxis experiment was made following Bargmann et al ^55^. On an NGM plate 1 μL of (-)-geosmin at the designated concentration was placed at 1 cm from one end of the plate, and on the other symmetrical opposite end, 1 μL of control was added (nanopure water). 1 μL of 1 M levamisole was added to each drop before drying them adjacent to a flame for 20-30min. Pure 2-butanone was used as a positive control ^56^. N2 adult *C. elegans* were washed 3 times in worm M9 buffer before addition at the centre of the plates (n=50). The plates were then incubated at room temperature for 1 hr before chemotaxis index calculation.

#### Behavioural assay

NGM agar with and without 0.54 μg/mL (-)-geosmin was poured into 24 well plates. Three adult *C. elegans* were added to each well and were video-taped for 10 min using a system that uses NIS Imaging BR version 3 software hooked up to a DSFi1c camera on a NIKON SMZ1500 microscope. The track line % 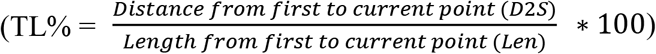 was calculated using Imaris 9.5 ^57^, data processed with a custom-built python script (File S6). Briefly, the script read the .xls files produced by Imaris particle tracking analysis, split the data into five two-minute segments, then extracted results for the track length and displacement of each worm for track line percentage determination. Videos were analysed using WormLab software 2020.1.1 ^58^ to generate the peristaltic speed (μm/s) and the head movement periodicity (μm) of the worms. Peristaltic speed is defined as peristaltic track length, the length of the track made by the worm during its movement, divided over time. Head movement periodicity is the wavelength of the sinusoidal wave created by tracing the head of the worms as they crawl. Worms that lodged in crevices and ceased moving were removed prior to analysis. Data was processed using Microsoft Excel 2011. Experiments were conducted in triplicate.

#### Predation assay

40μL of *S. coelicolor* (OD_600_ = 0.4-0.6) or *M. xanthus* (OD_600_ = 0.2-0.3) was added to the center of a 5 cm diameter NGM plate. The liquid was allowed to dry and the plates were then incubated at 30°C (7 days for *S. coelicolor* and 3 days for *M. xanthus*) Ten adult hermaphrodite *C. elegans* were then added at 2-5 mm away from the bacterial colony.

Quantification of the worms inside and outside of the bacteria was made at T= 0, 2, 4 and 24 hrs. Each experiment was conducted in at least triplicate.

#### Terpene add-in assay

Add-in experiments were made using *S. coelicolor* J2192, following the same procedure as in the predation assays with the following modification. One hour prior to the addition of *C. elegans* 40 μL of (-)-geosmin (2.25 μg/mL), (-)-2-methylisoborneol (40 μg/mL), or both was added to the bacterial colony and allowed to dry. Autoclaved nanopure H_2_O was used as a negative control. Triplicates of each added compound were performed with the addition of 10 adult hermaphrodite N2 *C. elegans* (n=30). Quantification was performed as per the predation assay.

#### *C. elegans* lethality assay

Solutions of (-)-geosmin (0.54, 5.4, 54 μg/mL) and 2-MIB (1.5, 15, 150 μg/mL) in LB broth were added to 24-well plates containing either NGM or NGM with *E. coli* OP50 as per H. Xiong et al. ^59^. LB broth was used as a negative control and a sodium hypochlorite solution (2.625 mg/mL) in LB broth was used as a positive control. The plates were incubated overnight at RT. Age synchronized plates of adult N2 worms were then washed in worm M9 buffer, dried on NGM plates, then added to each well (n = 5). Lack of response to touch stimuli was used to presume death. The number of dead *C. elegans* was quantified at T = 0, 2, 4 and 24 hrs. Experiments were run in triplicate.

#### Synthesis of 2-MIB

To a solution of (1R)-(+)-camphor (3.3 mmols) in anhydrous tetrahydrofuran (0.2 M) at 0 *°C* was added methylmagnesium bromide (4.95 mmols, 3 M). The reaction was then heated to reflux for 10 hours, before being quenched with a saturated solution of ammonium chloride in water. The THF was removed *in vacuo* and the remaining solution was extracted three times with EtOAc. The organic layers were then pooled and extracted with brine, then dried with sodium sulfate. The organic solvent was then removed *in vacuo* to give a white solid. (325.8 mg, 58.70 %). The crude mixture was then purified by flash chromatography (12% EtOAc in hexanes) to give the title compound as a white solid (48.7 mg, 8.77 % yield). ^1^H NMR (500 MHz, CDCl_3_) δ 2.07 (dt, 1H, *J* = 3.72 Hz, 13.08 Hz), 1.70 (m, 2H), 1.39 (m, 4H), 1.24 (s, 3H), 1.10 (s, 3H), 0.87 (s, 3H), 0.85 (s, 3H). ^13^C NMR (125 MHz, CDCl_3_) δ 79.6, 51.9, 48.9, 47.3, 45.4, 31.3, 27.0, 26.8, 21.4, 21.2, 9.9.

#### Statistical analysis

All statistical analyses were performed using student T-test calculations with two-tailed distribution and unequal variance. Statistical significance of *P* < 0.05 was noted as a red box in Figure 1D and as * for *P* < 0.05 or ** for *P* < 0.001 in Figure 2A-D. All replicates are distinct samples (biological replicates). Data was visualized with GraphPad Prism 9.0.

## Acknowledgements

We thank P. Prevost and the Forgione lab for assistance with the synthesis of 2-methylisoborneol; K. Flärdh for supplying *S. coelicolor* M145, J3003, and J2192; E. Despland for microscopy support; L.K. Freeman for discussions and feedback. Imaging support was provided by the Concordia Centre for Microscopy and Cellular Imaging, while CD and fluorimetry was performed with the assistance of the Concordia Integrated Platform for Biomolecular Function, Interactions and Structure.

## Author Contributions

BLF and LZ conceived the study. LZ performed all listed experiments with assistance from AI and YG (GC-MS); IO, KM and AP (C. elegans); CL (microscopy). IO and AP helped perform the behavioural, predation (S. coelicolor) and terpene add-in assays. KM and AP aided with the chemotaxis assay. CL prepared the python script. BLF and LZ prepared the manuscript, with feedback from all authors.

## Data and Material Availability

All data is available in the main text or the supplementary materials.

## Supporting Online Material

Supporting material is available online at BioRxiv.org

Materials and Methods

Figures S1 to S3

Tables S1 to S4

Files S1 to S6

