## Supplementary Information for "The ubiquitous terpene geosmin is a warning chemical"

Supplementary Material

### Authors

Liana Zaroubi[1], Imge Ozugergin[2], Karina Mastronardi[2], Anic Imfeld[1], Chris Law[2], Yves Gélinas[1], Alisa Piekny[2] & Brandon L. Findlay[1,*]

### Affiliations

[1]Department of Chemistry and Biochemistry, Concordia University, Montreal, Québec, Canada

[2]Department of Biology, Concordia University, Montreal, Québec, Canada

**This PDF file includes:**

Supplementary Text
Figs. S1 to S3
Tables S1 to S4

**Other Supplementary Materials for this manuscript include the following:**

Movies S1 to S5
File S1

### Supplementary Figures and Tables

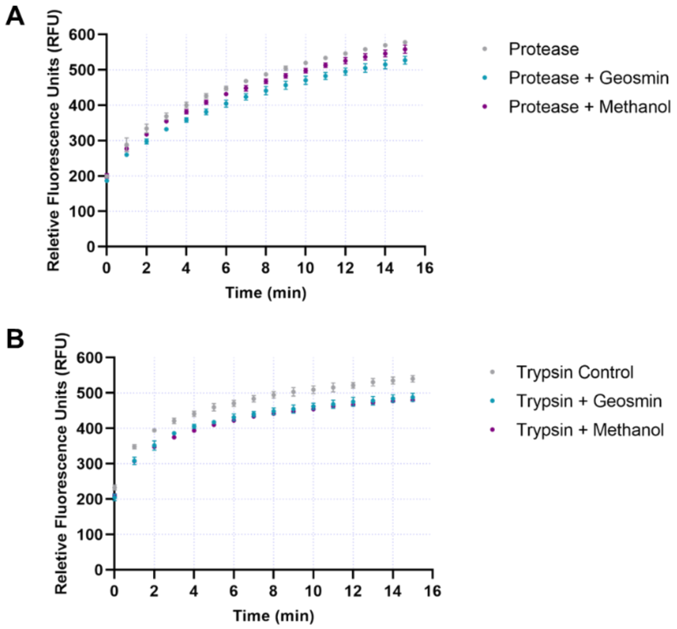


**Figure S1. Degradation of FTC-casein.** (A) Degradation of FTC-casein by *M. xanthus* DK1622 spent culture media. (B) Degradation of FTC-casein by trypsin. $\lambda_{excitation}$= 494 nm, $\lambda_{emission}$= 521 nm. Results of the protease control are shown in blue, protease with methanol in orange and protease with geosmin in grey.

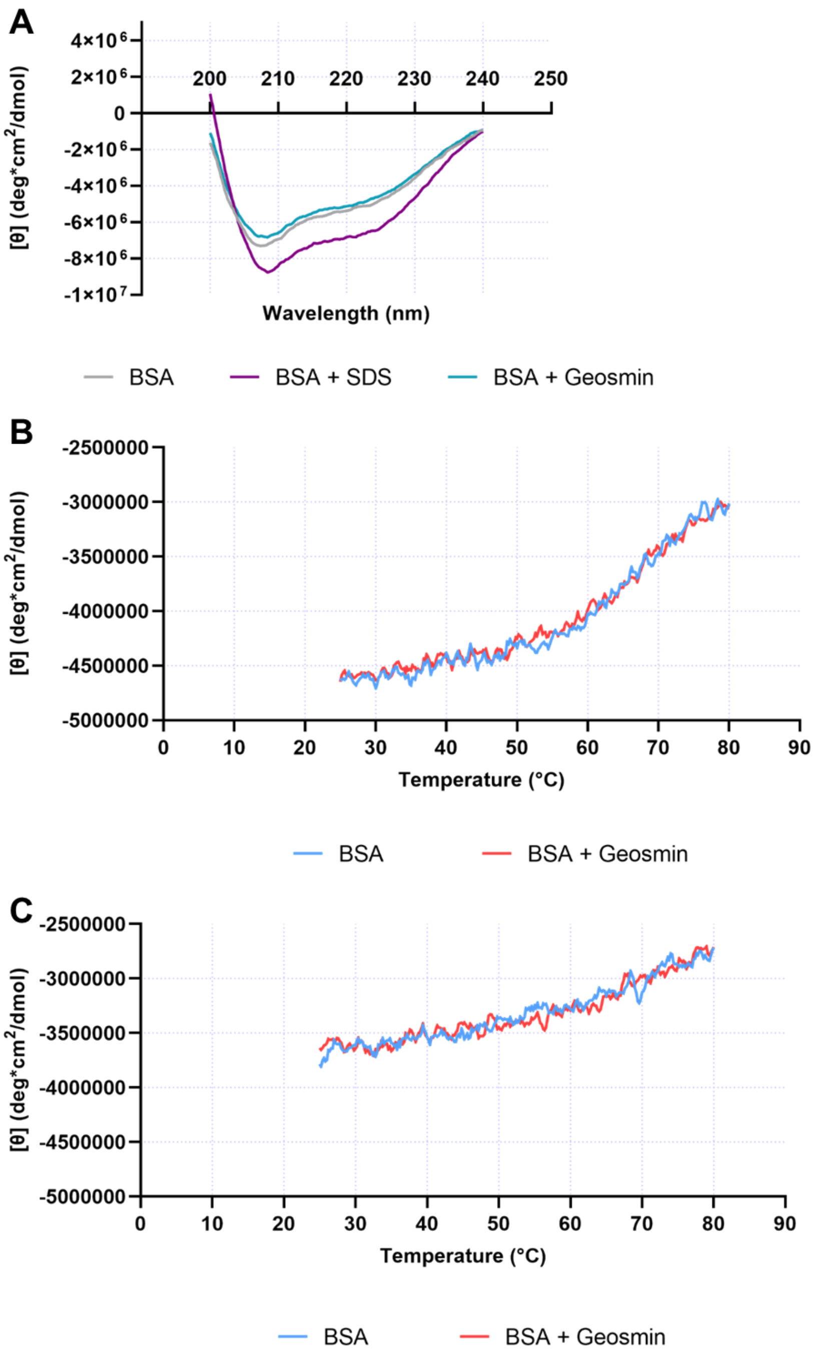


**Figure S2. Heat Denaturation of Bovine Serum Albumin (BSA).** (A) Heat denaturation of BSA at 80 °C. Denaturation of BSA during (B) heating and (C) cooling. The concentration of BSA is 1.25 µM, the concentration of geosmin is 0.18 mM, and the concentration of SDS is 0.75 mM. The wavelength window is from 200-400 nm.

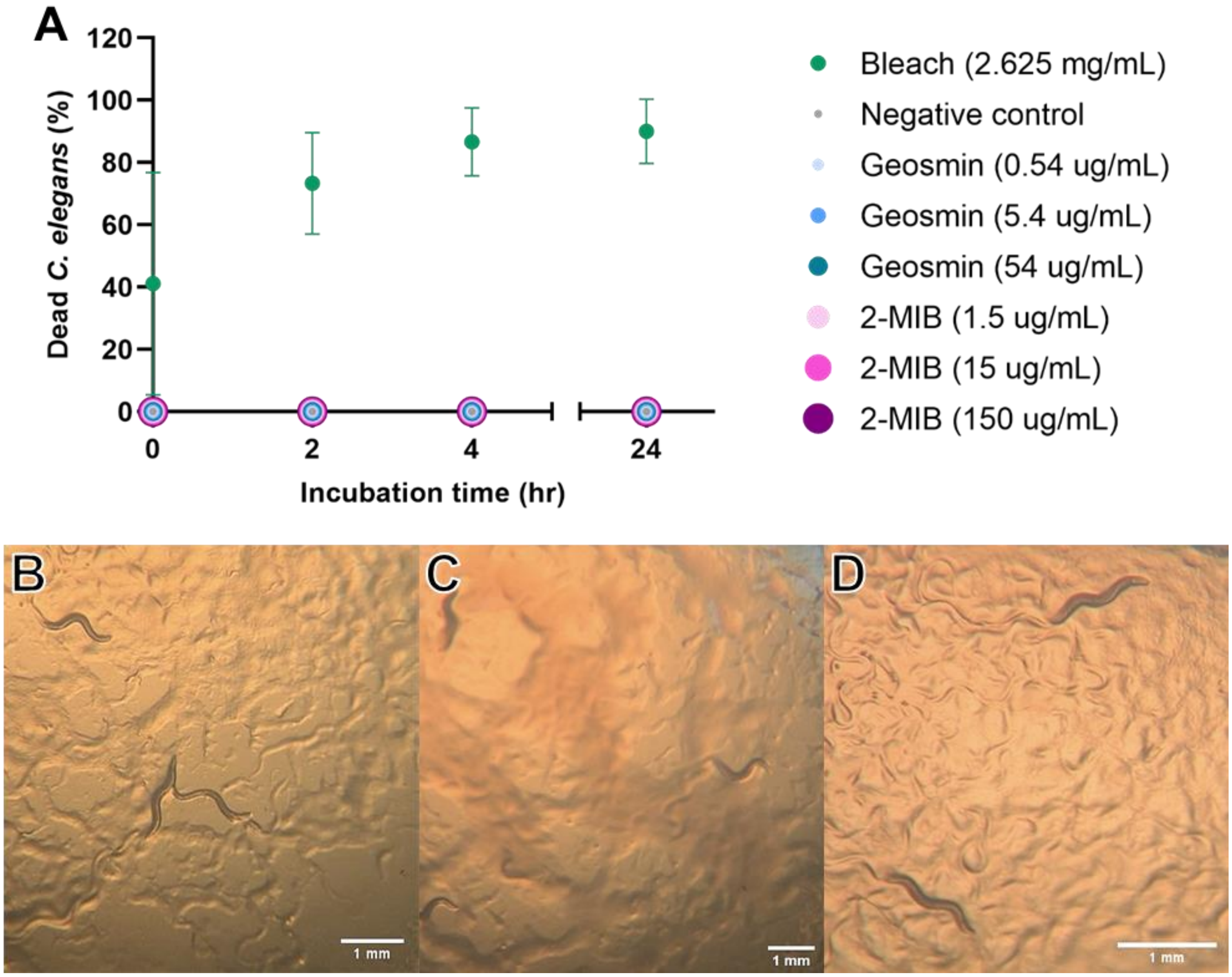


**Figure S3. Terpene toxicity and effect on *C. elegans* feeding. A,** *C. elegans* viability on a lawn of *E. coli* in the presence of geosmin, 2-methylisoborneol and bleach. **B**, Consumption of *E. coli* by *C. elegans*. **C**, Consumption of *E. coli* by *C. elegans* in the presence of geosmin (54 μg/mL). **D**, Consumption of *E. coli* by *C. elegans* in the presence of 2-methylisoborneol (150 μg/mL).

**Table S1: Quantification of cell-associated and extracellular geosmin.**

| Extraction Location | [Geosmin] (ppm) | |
|---|---|---|
| | Day 2.5 | Day 4 |
| Intracellular[a] | 0.015353 | 0.1151 |
| Extracellular[b] | 0.043 | - |
| Full[b] | 0.07 | 0.4105 |

[a]1:4 extraction with EtOAc.
[b]1:1 extraction with EtOAc.

**Table S2.** Minimum Inhibitory Concentration of Geosmin against Gram-negative and Gram-positive bacteria.

| Bacteria Strain | MIC (±)Geosmin (µg/mL) | MIC Methanol (µL)[a] |
|---|---|---|
| *M. luteus* DSM 20030 | 250 | 50 |
| *B. subtilis* DSM 10 | 250 | >50 |
| *B. thailandensis* E 264 | 62.5 | 6.25 |
| *K. aerogenes* ATCC 13048 | 500 | >50 |

[a] The geosmin sample was dissolved in methanol at a concentration of 2 mg/mL.

**Table S3: *C. elegans* N2 chemotaxis assay.**[a]

| [Geosmin] (μg/mL) | Worms in Geosmin Spot | Worms in Control Spot | Chemotaxis Index[b] |
|---|---|---|---|
| 54000 | 14 | 10 | 0.08 |
| 5400 | 15 | 7 | 0.16 |
| 540 | 10 | 10 | 0 |
| 54 | 10 | 5 | 0.1 |
| 5.4 | 6 | 12 | -0.12 |
| 0.54[c] | 4 | 14 | -0.2 |
| 0.054 | 17 | 8 | 0.18 |
| 0.0054 | 11 | 7 | 0.08 |
| 0.00054 | 11 | 5 | 0.12 |
| 2-butanone | 30 | 1 | 0.58 |

[a] Fifty adult *C. elegans* worms per trial. Singlicate experiments, unless otherwise noted.

[b] Chemotaxis index calculated as $\frac{Number\ of\ worms\ at\ geosmin\ spot - Number\ of\ worms\ at\ control\ spot}{Total\ number\ of\ worms}$.
The Average chemotaxis index is -0.1, s = 0.11.

[c] Experiment ran in triplicates.

**Table S4. Effect of geosmin on *C. elegans* movement.**

| Analysis | *C. elegans* | Mutant Deficiency | Control | | Geosmin | | *P*-value |
|---|---|---|---|---|---|---|---|
| | | | Average | Standard Deviation | Average | Standard Deviation | |
| **Track Line (%)** | N2 | Wild type | 0.4495 | 0.2464 | 0.5547 | 0.3115 | 0.0498 |
| | BR5514 | ADF, ASE, AWC | 0.4371 | 0.3032 | 0.5704 | 0.3091 | 0.0407 |
| | CE1248 | AWA | 0.5447 | 0.3098 | 0.3973 | 0.2477 | 0.0098 |
| | CX2065 | AWB, partial AWC | 0.5715 | 0.2969 | 0.7174 | 0.2617 | 0.0531 |
| | CX2205 | Olfactory system | 0.5676 | 0.2667 | 0.5015 | 0.3330 | 0.2376 |
| | CX5893 | AWC | 0.3797 | 0.2383 | 0.5124 | 0.3134 | 0.0233 |
| | NL2105 | Chemosensation | 0.5417 | 0.3210 | 0.5919 | 0.3253 | 0.4034 |
| | PR674 | ASE | 0.5617 | 0.2997 | 0.6174 | 0.2746 | 0.3523 |
| **Peristaltic Speed (µm/s) [Absolute peristaltic track length/time]** | N2 | Wild type | 30.30027 | 8.036627 | 40.89631 | 5.74027 | 1.13E-06 |
| | BR5514 | ADF, ASE, AWC | 28.60974 | 11.43549 | 28.62741 | 10.73547 | 0.8719 |
| | CE1248 | AWA | 38.47773 | 5.18592 | 30.76179 | 5.137974 | 1.55E-05 |
| | CX2065 | AWB, partial AWC | 31.69875 | 3.094122 | 45.48192 | 6.194929 | 5.57E-05 |
| | CX2205 | Olfactory system | 41.43203 | 1.540853 | 35.47248 | 1.379966 | 0.00353 |
| | CX5893 | AWC | 27.08556 | 9.214374 | 35.17561 | 5.317928 | 0.001547 |
| | NL2105 | Chemosensation | 24.34918 | 3.870071 | 26.03611 | 5.0001 | 0.2812 |
| | PR674 | ASE | 52.00835 | 3.823389 | 47.42004 | 6.401746 | 0.2098 |
| **Periodicity (µm)** | N2 | Wild type | 19.096 | 4.489433 | 19.49362 | 0.499788 | 0.03744 |
| | BR5514 | ADF, ASE, AWC | 18.12805 | 1.356552 | 17.20374 | 2.388581 | 0.2657 |
| | CE1248 | AWA | 16.38904 | 0.431564 | 12.12216 | 1.405526 | 2.44E-18 |
| | CX2065 | AWB, partial AWC | 16.08523 | 1.203601 | 18.68935 | 0.364582 | 1.94E-05 |
| | CX2205 | Olfactory system | 17.03812 | 0.239728 | 15.15458 | 0.604553 | 2.91E-05 |
| | CX5893 | AWC | 19.58694 | 4.840422 | 22.032 | 0.885555 | 0.09487 |
| | NL2105 | Chemosensation | 16.4984 | 2.778458 | 15.42235 | 0.691968 | 0.6440 |
| | PR674 | ASE | 22.65395 | 2.821125 | 21.9781 | 0.986451 | 0.1370 |

**Other Supplementary Files:**

**Movie S1.**
Video of *C. elegans* N2 crawling on agar.

**Movie S2.**
Video of *C. elegans* N2 crawling on agar that contains geosmin.

**Movie S3.**
Video of *C. elegans* NL2105 crawling on agar.

**Movie S4.**
Video of *C. elegans* NL2105 crawling on agar that contains geosmin.

**Movie S5.**
Nematode distress behaviour exhibited when in *S. coelicolor* colonies.

**File S1**
Imaris Python analysis script.
