## Supplementary material for "The ubiquitous terpene geosmin is a warning chemical": Supplmentary Files and Video: Supplementary File 1 - Python script.py

```
import numpy as np
import os,sys, io, string
import pandas as pd


#This section of the code makes a list of folders inside the directory at which you have pointed the code
fPath = sys.argv[1]
absFilePath = os.getcwd()
sampleList = os.listdir(fPath)
sampleList.remove('.DS_Store') #.DS_Store gets created by some computers when they see what's inside some folders; we don't need it, so we ignore it


e_length = 14 # this is the value to change to change epoch length

temp_length = np.floor(600/e_length)+1
temp_length = int(temp_length)
#I am pre-allocating arrays in which to store the data that will be calculated and saved at the end
data_out = np.empty((0,3),dtype=float) #arrays can be "empty"
epoch_Data = np.zeros((0,temp_length,5),dtype = float) #or you can initialise them as zeros
epoch_Data[:] = np.nan #this line changes every 0 in epoch_data (which is the whole array), into NaN values

#The line below initilises a list to which I can add values as we find new worms
worm_names = list()

#for every folder (sample) in the list of folders created above
for sample in sampleList:
    csvList = os.listdir(fPath+"/"+sample) #make a list of every csv file in the folder
    stem = csvList[0] #make a variable called stem, and set it as being equal to the name of the first csv file
    stem = stem[:-9] #remove the last 9 characters of stem, which will be the name of the .csv file, without _Area.csv
    #then we import the data from the Position file using pandas, and only import selected columns
    allData = pd.read_csv(fPath+"/"+sample+"/"+stem+"_Position.csv",header=2,usecols=(0,1,6,7,8)) #

    wormList = allData['TrackID'] #this gets the column containing the Worm IDs
    wormList = list(wormList)#This changes it from a Pandas dataframe to a python list
    wormList = set(wormList)#This changes it from a List, to a Set, which removes duplicate values

    #then, for every unique worm name in the set
    for worm in wormList:
        print(worm) #Show the worm name on the screen
        worm_names.append(stem+"_"+str(worm)) #add that worm name to the worm_names list we initialised earlier
        w_Straight = np.empty((1,0),dtype=float)#initialise somewhere to store the straightness data
        w_Distance = np.empty((1,0),dtype=float)#... and the distance data
        w_Displacement = np.empty((1,0),dtype=float)#...and displacement
        w_Speed = np.empty((1,0),dtype=float)#...and Speed
        wormData = allData.loc[allData['TrackID']== worm] #we use logical indexing in Pandas to pull out only the data that belongs to the worm we're looking at
        time = np.asarray(wormData['Time'],dtype = int) #...and isolate a new variable which is a list of timepoints in which the worm exists

        t0 = np.min(time) #the smallest value in this set is when the worm starts to exist
        tLast = np.max(time) #and the highest value is when it disappears
        tNow = t0 #set a temporary variable called tNow, and make it equal to t0 (when the worm appears)

        #I make a temporary array to store the data in; it has to have the same number of columns (21) as the array that will store the data from every other worm (epoch_Data, line 17)
        temp = np.zeros((1,temp_length,5),dtype = float) #make a zero array with 1 row, 21 columns and 4 layers to store data in
        temp[:] = np.nan #replace all 0s with NaNs

        e_tMin = int(np.floor(t0/e_length)) #this is the epoch in which the worm starts to exist
        e_tMax = int(np.ceil(tLast/e_length)) #column in which the worm stops existing
        for epoch in range(e_tMin,e_tMax): #for every epoch in which the worm exists

            #isolate the data that corresponds to this epoch
            timeSlice = wormData.loc[wormData['Time']<=(epoch*e_length)+e_length]
            timeSlice = timeSlice.loc[timeSlice['Time']>=(epoch*e_length)+1]

            #get the X and Y values
            x_pos = np.asarray(timeSlice['Position X'],dtype = float)
            y_pos = np.asarray(timeSlice['Position Y'],dtype = float)

            #if len(x_pos) == e_length: #If the worm exists for the full 20s
            x0 = x_pos[0] #Where does it start this epoch
            y0 = y_pos[0]
            x1 = x_pos[-1] #Where does it finish this epoch ([-1] means the last item in an array)
            y1 = y_pos[-1]
            displacement = np.sqrt((x1-x0)**2+((y1-y0)**2))#This is the distance between XY1 and XY0
            distance = 0 #We then start counting distance (at 0, of course)
            t = 1 #record how many seconds the worm existed in the epoch
            for timePoint in range(len(x_pos)-1): #for every timepoint a worm exists, except the last
                x0 = x_pos[timePoint] #where does it start
                y0 = y_pos[timePoint]
                x1 = x_pos[timePoint+1]#where does it go in the next timepoint
                y1 = y_pos[timePoint+1]
                distance = distance + np.sqrt((x1-x0)**2+((y1-y0)**2)) #add that value to the total distance travelled
                t = t + 1#this worm existed for another second - at the end of all but the first and last epoch, t should equal 20

            speed = distance / t #how fast does it move?
            straightness = displacement/distance #how straght was the track?
            temp[0,epoch+1,0] = straightness #store these calculated values in separate layers of the temporary array initialised above
            temp[0,epoch+1,1] = distance
            temp[0,epoch+1,2] = displacement
            temp[0,epoch+1,3] = speed
            temp[0,epoch+1,4] = t

        epoch_Data = np.append(epoch_Data,temp,axis=0) #append the values from this worm onto the big array contain all worm data

print(len(worm_names)) #These are just some outputs to check it's reading everythingin correctly
print(epoch_Data.shape)

#We then make a pandas dataframe for each statistic we calculated, and make sure they have a column for the worm Names
epoch_straightness = pd.DataFrame(data = epoch_Data[:,2:,0]) #pull the 0th layer for straightness
epoch_straightness['Names'] = worm_names
epoch_distance = pd.DataFrame(data = epoch_Data[:,2:,1])#pull the 1st layer for distance
epoch_distance['Names'] = worm_names
epoch_displacement = pd.DataFrame(data = epoch_Data[:,2:,2])#pull the 2nd layer for displacement
epoch_displacement['Names'] = worm_names
epoch_speed = pd.DataFrame(data = epoch_Data[:,2:,3])#pull the 3rd layer for speed
epoch_speed['Names'] = worm_names
epoch_tCount = pd.DataFrame(data = epoch_Data[:,2:,4])#pull the 4th layer for timepoint count
epoch_tCount['Names'] = worm_names

#save these dataframes to csv files, and enjoy looking at them!
epoch_straightness.to_csv(fPath+"/epoch_straightness.csv")
epoch_distance.to_csv(fPath+"/epoch_distance.csv")
epoch_displacement.to_csv(fPath+"/epoch_displacement.csv")
epoch_speed.to_csv(fPath+"/epoch_speed.csv")
epoch_tCount.to_csv(fPath+"/epoch_time.csv")
```
